# Neural population dynamics of human working memory

**DOI:** 10.1101/2022.09.23.509245

**Authors:** Hsin-Hung Li, Clayton E. Curtis

## Abstract

Temporally evolving neural processes maintain working memory (WM) representations of information no longer available in the environment. Nonetheless, the dynamics of WM remain largely unexplored in the human cortex. With fMRI, we found evidence of both stable and dynamic WM representations in human cortex during a memory-guided saccade task. The stability of WM varied across brain regions with early visual cortex exhibiting the strongest dynamics. Leveraging population receptive field modeling, we visualized and made the neural dynamics interpretable. Early in the trial, neural responses in V1 were dominated by narrowly tuned activation at the location of the peripheral target. Over time, activity spread toward foveal locations and targets were represented by diffuse activation among voxels with receptive fields along a line between the fovea and the target. We suggest that the WM dynamics in early visual cortex reflects a transformation of sensory inputs into abstract task-related representations.

---

Working memory (WM) allows us to store information over a brief period of time, increasing the duration of neural representations available to support computations and decisions. Early studies on the neural mechanisms of WM focused on the prefrontal cortex (PFC) of non-human primates^1,2^, reviewed in^3^, and demonstrated that neurons in PFC exhibit elevated and sustained activity during memory delays suggesting a mechanism by which stimulus information is maintained^4–11^.

Implicit in the studies of persistent activity is the idea that the population neural response that encodes a specific feature value is stable through memory delay. However, later empirical studies reported that during memory delays, some neurons in PFC exhibit temporal dynamics more complicated than those predicted by stable persistent activity^12–16^. Recent multivariate analyses revealed that both stable and dynamic codes of WM coexist in the macaque PFC at the population level^17,18^. Fewer studies have investigated the dynamics of WM in the human brain. By measuring the responses evoked by task-irrelevant stimuli presented during memory delay, previous human EEG studies inferred that the neural code of WM exhibited dynamics 19,20

Here we address several open issues regarding the neural code of WM and its dynamics in the human brain. First, going beyond a single region of interest and leveraging the findings that the content of WM can be decoded from fMRI voxel activity patterns across widespread brain regions^21–43^, we quantified and compared the stability of WM codes across multiple brain regions. Second, we aim to elucidate the driving factors of WM dynamics by projecting population neural response into the visual field by population receptive field (pRF) mapping^44,45^, making the neural dynamics of WM interpretable.

To preview, we found that stable and dynamic codes coexist in most of the brain regions, and most importantly, the stability of WM varied across brain regions with the early visual cortex exhibiting the strongest dynamics while the high-level visual and the parietal cortex showed higher stability. In V1, where we observed prominent dynamics, neural activation was limited to the location of the peripheral target early in the delay, and over time, spread toward the foveal visual field along the polar angle of the target. These results demonstrate how WM dynamics can be driven by different temporal dynamics among neural populations with different selectivity (eccentricity in this case) and may reflect a recoding of sensory inputs into a format most proximal to the demands of the task.

## Results

To facilitate direct comparisons with the existing monkey neurophysiological studies eg^4,6,9,17,18,46^, we used a memory-guided saccade task to study spatial WM. In each trial, a brief (500 ms) target dot was presented in the periphery, followed by a 12 s delay period. The polar angle of the target was chosen pseudo-randomly from 1 of 32 positions spanning the full circle. After the delay, participants reported the remembered location with a memory-guided saccade (Fig. 1A). Participants were able to make precise memory reports close to the target locations (Fig. 1B). In addition to the memory-guided saccade experiment, participants underwent a pRF mapping session^44,45^, which allowed us to define four retinotopic visual (V1, V2, V3, and V3AB) and four parietal (IPS0, IPS1, IPS2, and IPS3) areas as the regions of interest (ROIs).

**Figure 1.**
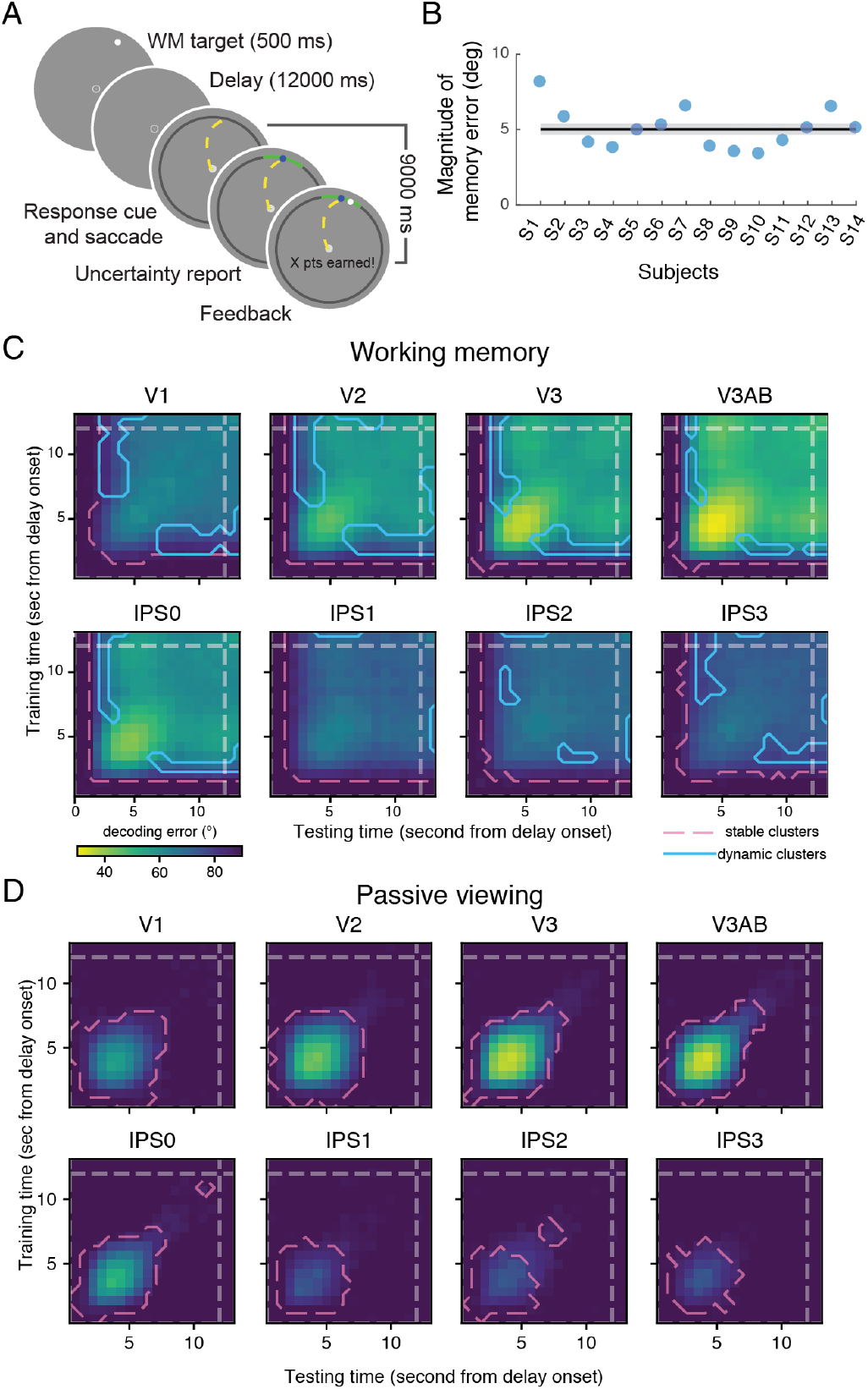
Behaviors and temporal generalization analysis. (A) Procedures. Participants maintained fixation while remembering the location of a target dot presented at a pseudorandom polar angle and at 12° eccentricity from fixation. At the end of each trial, participants made saccadic eye movements to report their memory, and adjusted the length of an arc for uncertainty report. (B) The mean magnitude (absolute value) of memory error of each participant. The horizontal line represents the group mean with ±1 s.e.m. (C) The temporal generalization analysis. Decoders were trained to decode the target location from the fMRI BOLD response. The temporal generalizability of the decoders trained with voxel activity patterns of each time point were tested with the data of all the time points. Pink dashed lines: the stable clusters, the cluster that exhibited above-chance decoding performance. Blue solid lines: the dynamic clusters, the off-diagonal elements that exhibited lower decoding performance than (both of) their corresponding diagonal elements. (D) Same as (C) but for the passive viewing experiment. Only stable clusters were observed.

### Coexistence of stable and dynamic WM codes

We first characterized the dynamics of the WM neural code by a temporal generalization analysis^18,47,48^, in which we tested whether the decoder trained by the voxel activity pattern at one time point can be generalized to decode the location (polar angle) of the WM target using the voxel activity at other time points. We found a stable WM code in all the ROIs (pink dashed lines in Fig. 1C), quantified by the above-chance decoding performance among the off-diagonal elements of the cross-decoding matrices. That is, the location of the WM target remained decodable even when the training data and the testing data came from different time points. In addition, across all the ROIs, WM content was present shortly (about 1 to 2 sec) after the delay onset (or target offset), and remained decodable throughout the delay period. We next asked if WM representations changed over time by testing if the off-diagonal elements showed worse decoding performance than their corresponding diagonal elements. In most ROIs (V1, V2, V3, V3AB, IPS0, IPS2 and IPS3), we observed clusters of off-diagonal elements exhibiting reduced decoding performance, indicating a dynamic WM neural code (light blue lines in Fig. 1C), which coexisted with the stable neural code.

Note that the decodable signals we observed throughout the delay do not merely reflect a slow decay of sensory-evoked response, but rely on WM maintenance. A subset of the subjects (n=6) participated in a passive viewing experiment in which they were asked to perform a task at central fixation while a high-contrast flickering peripheral ‘WM target’, treated as an irrelevant stimulus, was present continuously throughout the delay. In this case, we only observed a stable code without significant dynamics, and the peripheral ‘WM target’ was only decodable for a much shorter period of time early in the delay without sustaining through the delay (Fig. 1D).

### Neural subspaces

We used principal component analysis (PCA) to visualize the neural subspaces that best explain the variance of voxel activity across different target locations. As an approach complementary to temporal generalization, PCA allows us to further (1) inspect whether target locations are encoded topologically, reflecting their relationships in the visual field space (2) investigate the geometry underlying the dynamics (3) quantify and thus compare the stability across brian regions.

The existence of a stable code indicates a coding subspace within which target locations are stable and maintain their relative positions throughout the delay. We characterized such a stable subspace by applying PCA on time-averaged BOLD responses, discarding the dynamical aspect of the data^17^. We focused on a subspace defined by the top two principal components that maximized the variance explained. We projected the data of each single time point during the delay into this stable subspace and found that in this subspace, target locations were topologically organized, preserving their spatial relationships in the visual field, and the neural activity remained stable across time in this subspace (Fig. 2).

**Figure 2.**
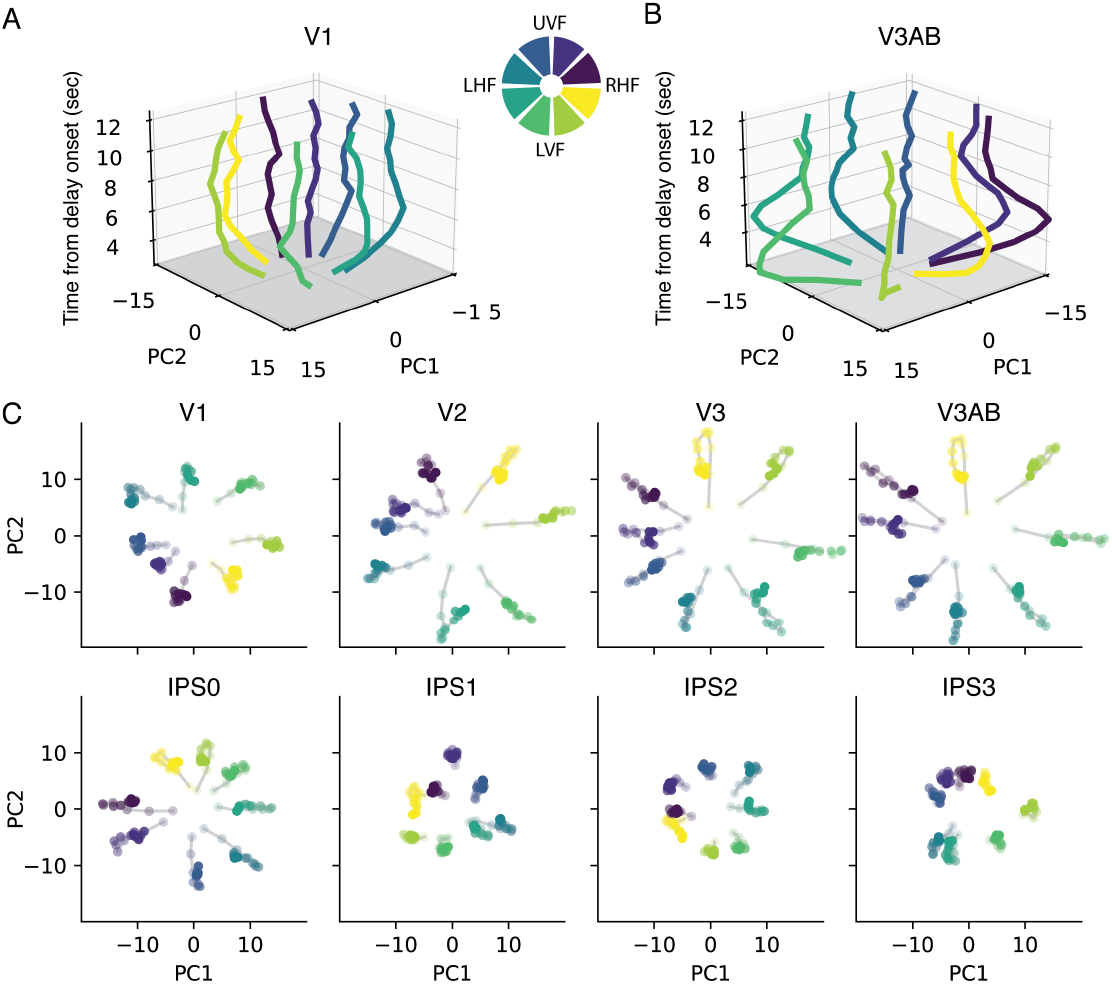
Stable subspaces. The stable coding subspaces were identified by applying PCA to time-averaged voxel activity patterns. (A) V1’s response at each time point during the delay period was projected into the stable subspace, the top two principal components (x-axis and y-axis) obtained from PCA. The z-axis represents time from delay onset. Each curve represents one bin of target location. The legend indicates the location, in the visual field, represented by each of the eight colors. (B) Same as (A) but for V3AB. (C) Similar to (A) and (B) but without visualizing the time axis (z-axis); thereby is equivalent to the bird-eye’s view of (A) and (B). Each dot represents data from one time point during the delay, with more saturated colors representing later time points.

We next investigate the dynamical aspect of the neural code by dividing the 12-second delay into three—the early, middle and late—time windows with equal length and computing their corresponding neural subspaces. A potential source of the dynamic code is rotational dynamics where targets move in a subspace across time but maintaining their relative positions^49–51^. We did not observe such rotations when projecting data of individual time points (Fig. 3A), or of each time window (Fig. 3B; also see Supplementary Fig. 1), into the early, middle or late subspace. We did observe that the variance of data was best explained by the neural subspace derived at the same time points. Overall, these results indicate that the WM dynamics is driven by changes of neural subspace where different neural populations were used to represent the targets at different time points (Fig. 3D).

**Figure 3.**
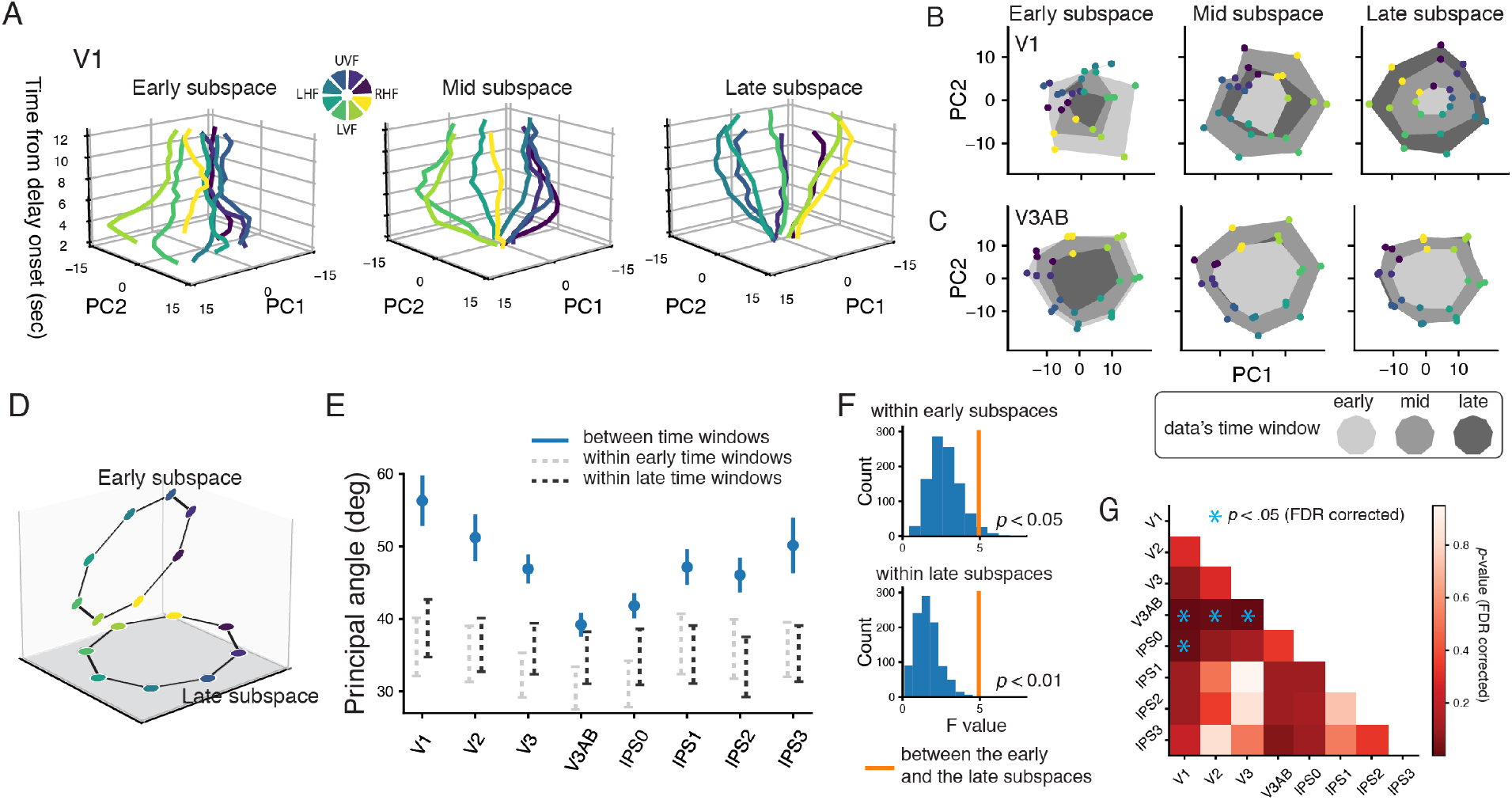
Dynamic subspaces. Early, middle and late neural coding subspaces were identified by applying PCA to voxel activity patterns in three different time windows. (A) V1’s response of each time point during the delay was projected into the early, mid and late subspaces, where the top two principal components (x-axis and y-axis) were derived from PCA. The z-axis represents the time from delay onset. Each curve represents one bin of the target locations. (B) Projections of the early, mid and late (data points connected with areas with three different gray levels) voxel activity patterns of V1 into the early, mid or late subspaces (from the late to right panel). (C) Same as (B) but for V3AB. See Supplementary Fig. 1 for the projections of other ROIs. (D) Cartoon illustration of two subspaces from different time points that encodes the WM target locations in the high dimensional space. (E) The dark blue data points represent the principal angle, between the early and late subspaces, of each ROI (group mean with ±1 s.e.m.). The dashed lines represent 95% confidence intervals of the distribution of the principal angles computed between the two subspaces estimated by resampling the data from either the same early (light gray dashed lines) or the late (dark gray dashed lines) time windows. (F) The main effect of ROI on the principal angle between the late and the early subspaces is larger than that predicted by the principal angle within the same time window. Top: by a bootstrapping procedure, in each iteration, we resampled the data, estimated two early subspaces and computed the principal angle between them. We then computed the main effect of ROI on the principal angle by ANOVA, leading to a bootstrapped distribution of the F values (blue histograms). P value was computed by comparing the bootstrapped distributions with the empirical F value computed using the principal angle between the early and the late subspaces (orange vertical lines). Bottom: same as the top figure, but the bootstrapped distribution was computed by resampling the late subspaces. (G) Pairwise comparisons of the principal angles between ROIs.

To compare the stability of neural code across brain regions, for each ROI, we quantified how much the neural subspace changed across time by the principal angle between the subspaces of the early and the late time windows (see Methods)^52–54^. We found that the stability of WM representations, quantified by principal angles, varied across ROIs (*F*(7, 91) = 3.88, p < .00). WM representations underwent greater changes in early visual cortex, and were more stable in higher-level visual cortex and parietal cortex. (confirmed by pairwise comparisons in Fig. 3G). We obtained similar results when using the ratio of variance explained (Supplementary Fig. 2). We also computed the principal angles between two subspaces that were estimated by the data resampled from the same time window (dashed lines in Fig. 3E). As expected, these baseline values were smaller than the angle computed between time windows, and more importantly, they did not show variation across ROIs like what we observed for the principal angles computed across time (Fig 3F). Thus, the stability of WM differed across ROIs, which cannot be explained by factors such as the reliability of subspace estimation.

### Factors driving WM dynamics

To understand and interpret the WM dynamics we observed, we utilized the voxels’ pRF (center and size) to compute activation maps by projecting voxel activity patterns into the coordinates of two-dimensional visual field space (Methods). Here, we focus on two ROIs—V1 and V3AB—where we observed the strongest and the weakest dynamics (see Supplementary Fig. 3 to 8 for other ROIs).

In V1, the spatial pattern of the population neural response showed clear changes across time. The response first emerged at the target’s polar angle and eccentricity, consistent with previous fMRI studies that visualized neural responses in the visual field space^35,43,55–57^. After about 4.5 seconds, the response at the target’s eccentricity declined, and spread across eccentricity, spanning the visual field between the target and the fovea (Fig. 4A and 4B). Toward the end of the delay, activity peaked at the foveal region with a tail pointing toward the target. We observed sequential activity when binning the locations on the activation maps (at target’s polar angle) by eccentricity—response at the target’s (far) eccentricity exhibited the earliest peak, followed by intermediate locations, while the foveal locations peaked at the latest time points (Fig. 4C). Overall, the neural representations for the WM target changed from a bump at the target into a more diffuse line-like pattern over time (Fig. 4B). In V3AB, the ROI with the greatest stability, we found that the peak of activation remained at the target’s peripheral location over the course of the trial (Fig. 5). Notably, when visualizing the activation maps for V1 during the passive viewing experiment, we did not observe dynamics like those in the WM experiment. The response emerged at the target’s polar and eccentricity at the early time points, and diminished in the middle of the delay (Fig. 6).

**Figure 4.**
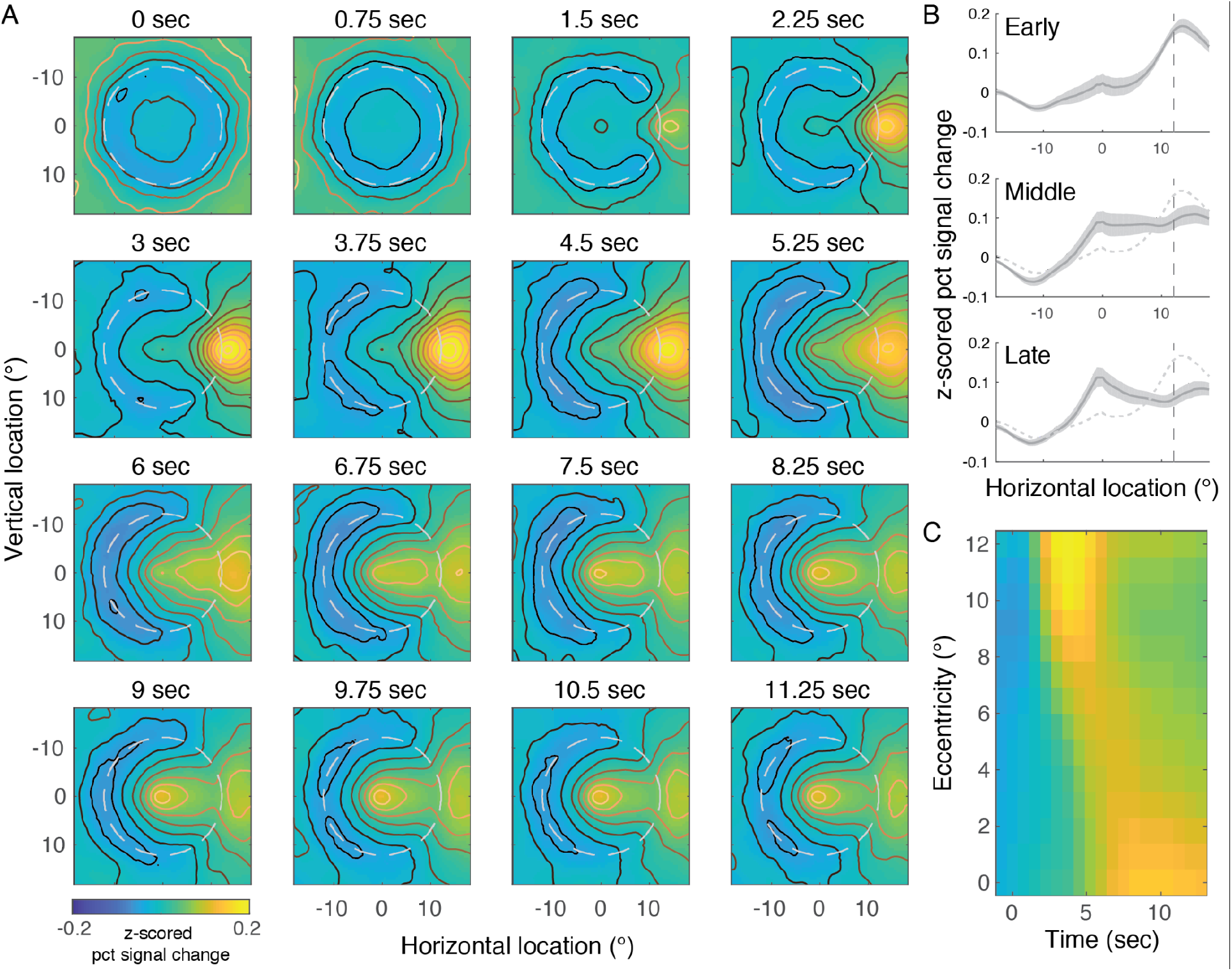
Visualization of WM dynamics in V1. (A) Activation maps visualize the projection of voxel activity patterns onto two dimensional visual field space. Each image represents the activation map reconstructed for each TR during the delay. (B) The horizontal slice over the activation map at vertical location = 0° for each time window (5 TRs per time window. TR at 0 sec was not included). For comparisons, the activity of the early time window was plotted as dashed curves in the middle and late time windows. Data represents mean ± s.e.m. (C) Population response for different eccentricities as a function of time, computed by averaging over the activation maps constrained within a sector spanning ± 30° centered the target’s polar angle and segmenting the sector into bins from far to near eccentricity in steps of 1°.

**Figure 5.**
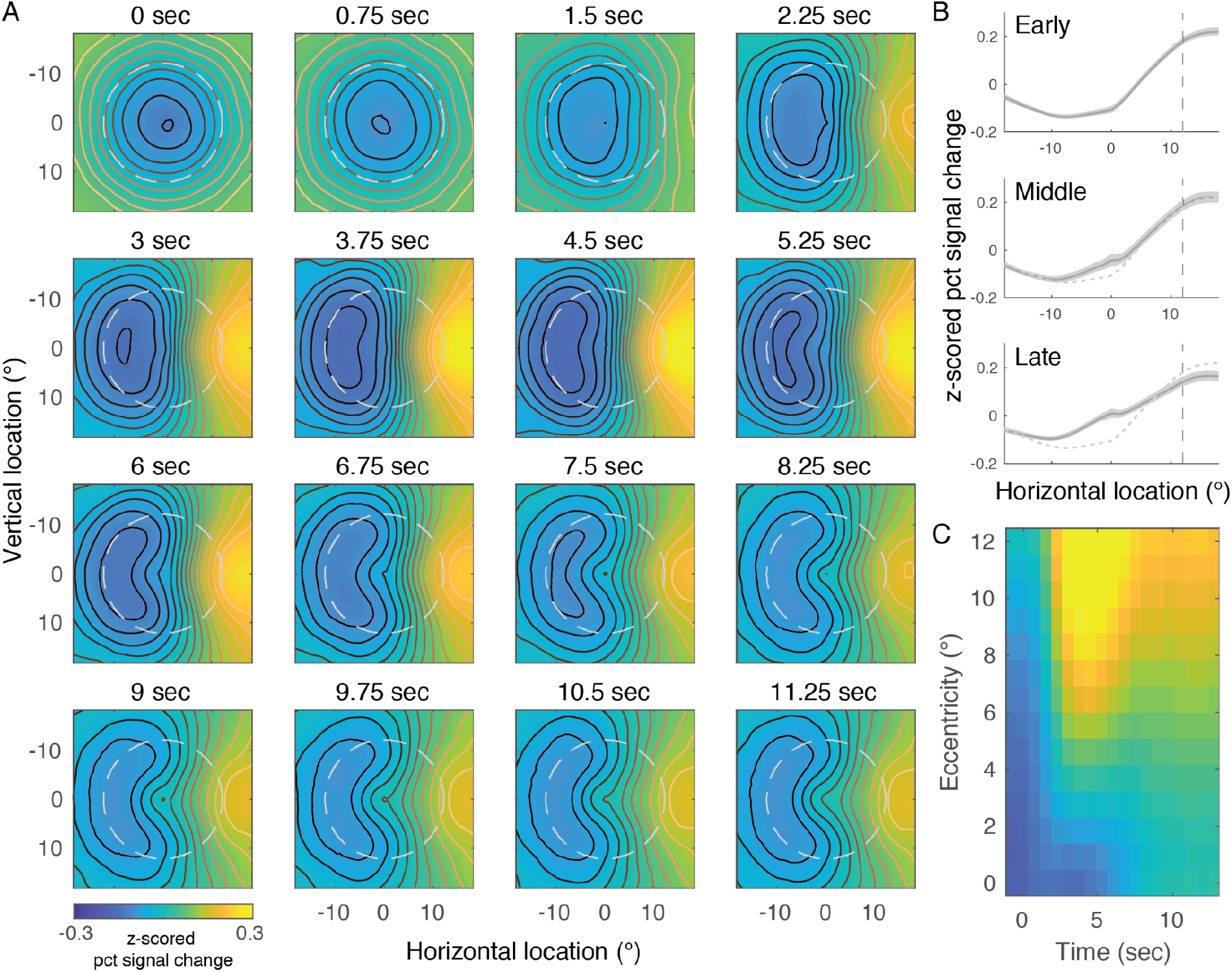
Visualization of WM dynamics in V3AB. (A) to (C) plotted in the same format as those in Figure 4A to 4C, but for V3AB.

**Figure 6.**
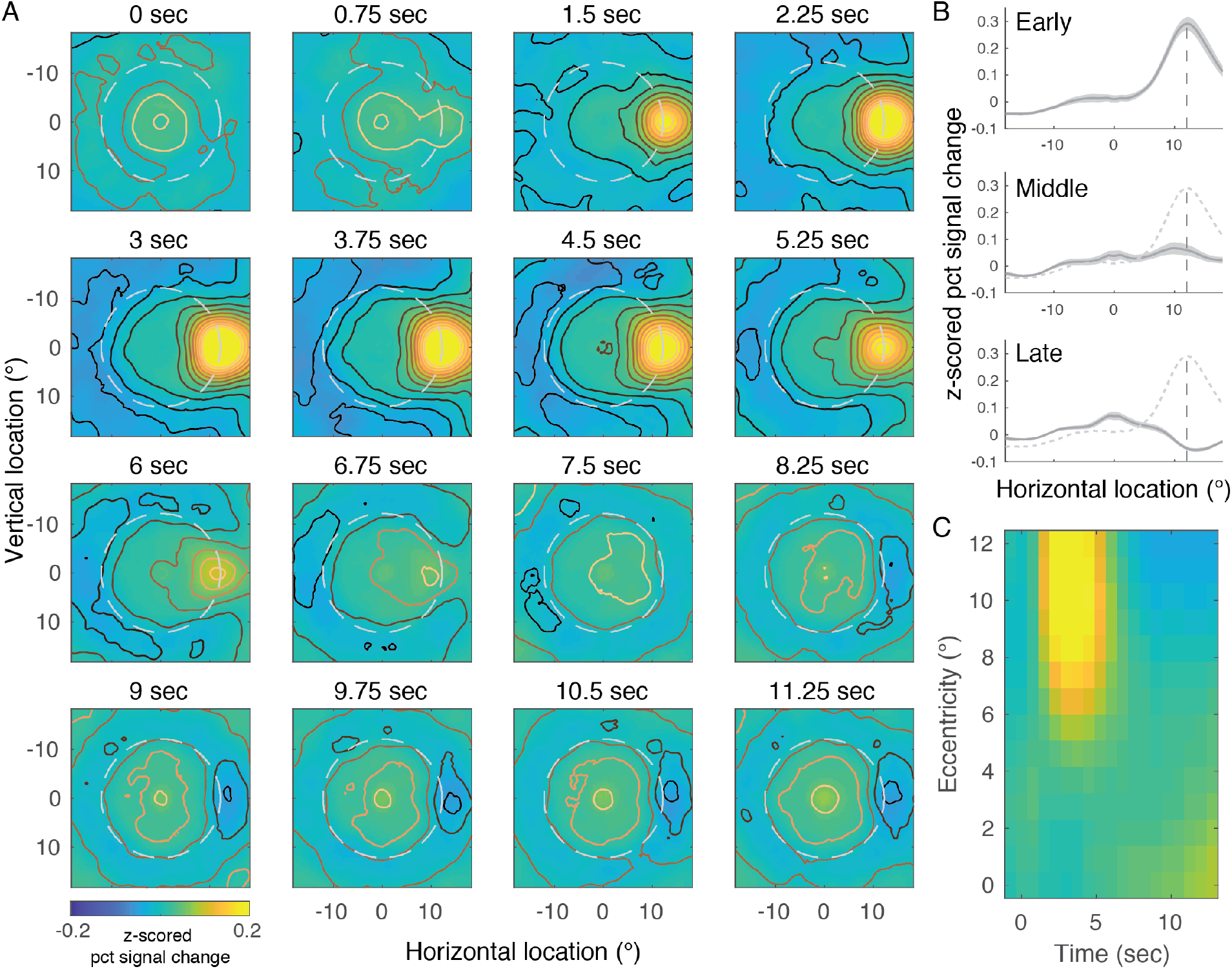
Visualization of neural dynamics in V1 during passive viewing experiment. (A) to (C) plotted in the same format as those in Figure 4A to 4C, but for the passive viewing experiment.

Besides eccentricity, changes of neural population’s selectivity for polar angle could also contribute to WM dynamics. We computed polar angle response functions by collapsing the two dimensional activation maps across eccentricity (Fig. 7A and 7B). We fitted von-Mises functions to the response at each time point with the center, gain and width as the free parameters. All ROIs had response functions that peaked around the target’s polar angle. For both V1 and V3AB, the gain of the response function peaked at about 4-5 seconds and remained above zero throughout the delay. The width of the functions showed different dynamics between the two ROIs (Fig. 7E). In V3AB, the width remained stable once it reached its lowest level (similar patterns were observed in all the ROIs in IPS, Supplementary Fig. 9). In contrast, the width in V1 first reached a lower value than V3AB, which is expected as V1 has a smaller pRFs when measured by retinotopic mapping procedures^45,58^, but increased afterward and became as wide as the width in V3AB. This dynamics may indicate that the neural activity observed in V1 reflects feedback signals during the later time points during the delay (see Discussions).

**Figure 7.**
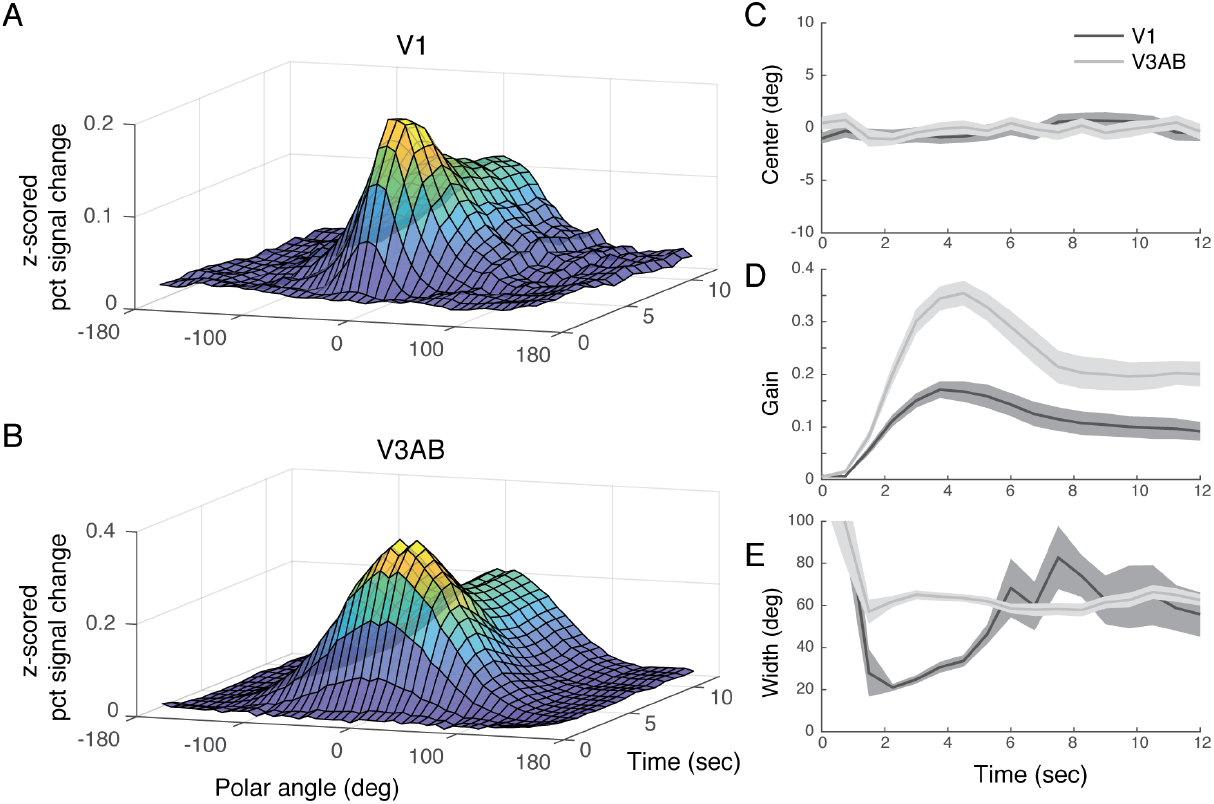
Polar angle response functions. (A) V1’s polar angle response functions as a function of time from delay onset. (B) Same as (A), but for V3AB. (C) The center of the response functions. (D) The gain of the response functions. (E) The width of the response function. Data represents mean ± s.e.m.

### Neural code during WM is stable in PFC

Previous studies of the neural population codes during WM in nonhuman primates have largely focused on the neural activity in PFC. Here, we extended our analyses above to two frontal regions, iPCS and sPCS, the regions where we previously observed topographic organization in the pRF mapping session consistent with^45^ We found decodable WM content in both frontal regions but their decoding errors were larger than all the other ROIs (Fig. 8A). In the temporal generalization analysis, we found a stable cluster throughout the memory delay in each of the ROIs, without significant dynamical clusters (Fig. 8B and 8C). Projecting the voxel activity pattern of each time point into the stable subspace extracted by PCA, we found that the locations of the targets remained largely stable and separable within the stable subspace (Fig. 8D and 8E). However, their spatial topology (in the visual field space) was not well-maintained like those observed in the visual and the parietal cortices.

**Figure 8.**
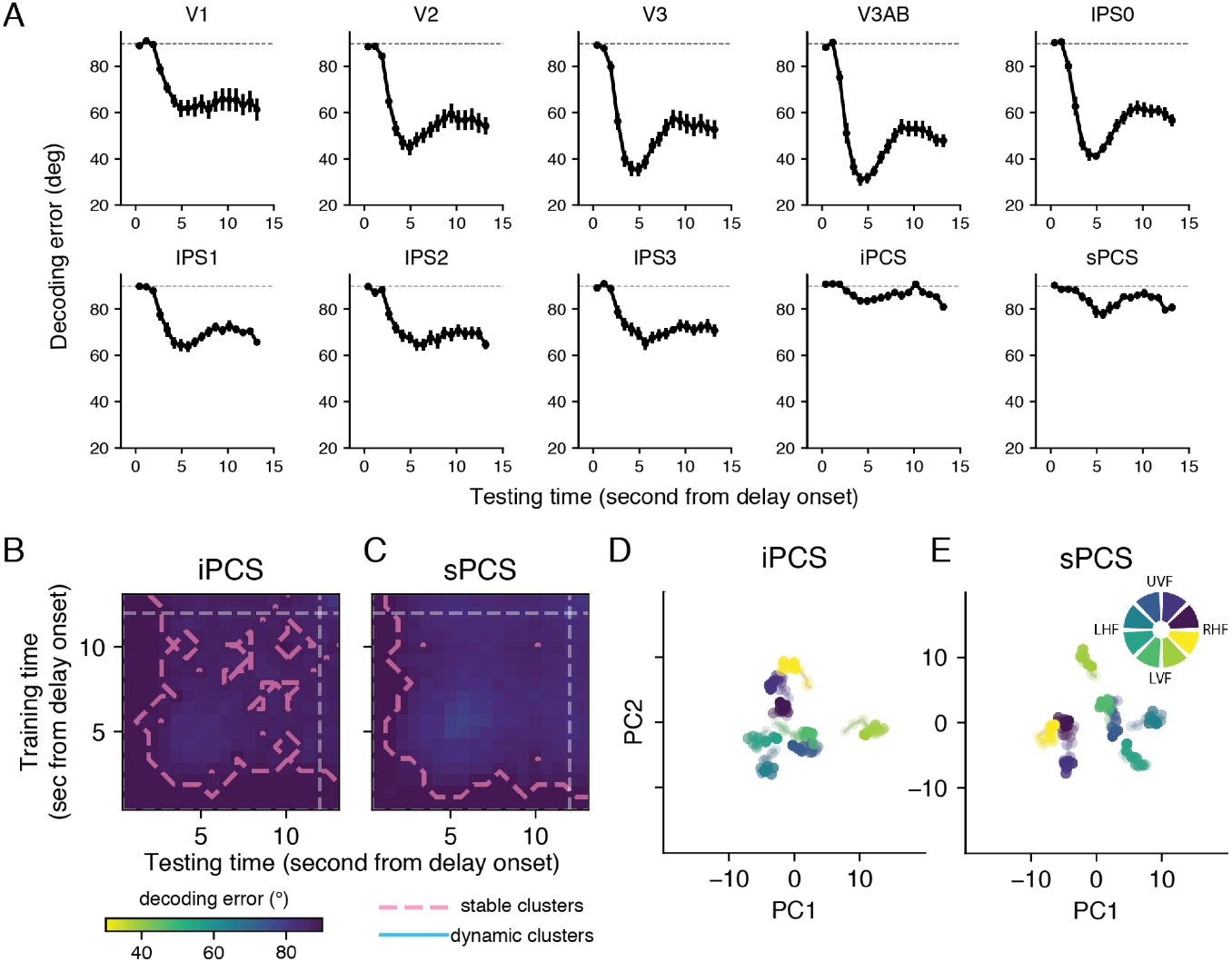
WM decoding in PFC. (A) Decoding error of each ROI including two ROIs in the prefrontal cortex. These data are equivalent to the diagonal elements of the decoding matrices from the temporal generalization analysis (Figure 1C and 8B-C). Data points represent group mean with ±1 s.e.m. (B-C) Temporal generalization analysis for iPCS and sPCS. Plotted in the same format as Figure 1C. (D-E) Stable subspaces. Plotted in the same format as Figure 2C. The stable coding subspaces were identified by applying PCA to time-averaged voxel activity patterns. Voxel activity pattern of each time point during the delay period was projected in the stable subspace, the top two principal components (x-axis and y-axis) obtained from PCA.

## DISCUSSION

To summarize, in the human brain, we found evidence of both stable and dynamic WM representations in a simple memory-guided saccade task (Fig. 1). By dimension reduction and analyzing the geometry of neural subspaces, we showed that the dynamics of WM varied across brain regions, with the early visual cortex exhibiting the greatest changes across time (Fig. 3). Projections of the voxel activity on visual field space revealed that the dynamics of WM in early visual cortex involves a propagation from neural populations selective for the peripheral location of targets to locations near the fovea (Fig. 4), accompanied by a decrement in polar angle selectivity (Fig. 7).

The first part of our results—temporal generalization and PCA—show striking similarities, but also differences, to those observed in neurophysiological recordings in monkey PFC. Applying the same temporal generalization analysis to data from Watanabe and Funahashi^59^, Spaak et al.^18^also reported the coexistence of stable and dynamic coding in monkey’s PFC for spatial WM. In both ours and their results^18^, in the cross-decoding matrices, the dynamic code exhibited as a horizontal elongated cluster (and a vertical mirrored version). Similarly, in a reanalysis of Constantinidis et al.,^60^and Romo et al.,^8^, Murray et al.^17^reported that a stable subspace explained a comparable amount of stimulus variance relative to a dynamic subspace during the delay. Only around the time of stimulus presentation did the stable subspace show a disadvantage compared to the dynamic subspace. Together, these results underscore that the strongest dynamics of WM occurred during the transition between a brief perceptual encoding phase and a longer mnemonic maintenance phase, and they are in line with the notion that mnemonic codes are somehow different from perceptual codes given that classifiers trained on fMRI patterns during visual stimulation often showed lower decoding performance during WM maintenance^22,42,43^. Moreover, WM representations remain decodable in the presence of visual distractors^37,40,42^ suggesting that because these representations do not cause catastrophic interference, they perhaps have unique representational formats^61^.

Despite the similarities between our findings and the neurophysiological results from monkey PFC regarding the information at the population level, we find evidence for coexisting stable and dynamic WM codes in early visual cortex, not PFC. While we cannot know whether these differences are due to differences in species or differences in measurements of neural activity, they parallel a vast literature in humans demonstrating the importance of early visual cortex for WM^3^.

Our results revealed that even under the simplest spatial WM task, different brain regions represent WM content in different formats. Neural response in higher-level visual cortex and parietal cortex almost exclusively represented the memorized locations, peaking at the target’s polar angle and eccentricity throughout the delay. In contrast, the early visual cortex seems to concurrently reflect the immediate task demand that requires observers to concurrently hold fixation and the memorized target at far eccentricity, leading to an activity with a peak at more foveal locations with a tail pointing toward the target. Additionally, the activation we observed in early visual cortex during the late time points of the delay might represent the planned trajectory, or the rehearsal, of the memory-guided saccade. In this case, the sensory properties of the visual target were reformatted into a mnemonic code more proximal to the response required by the task. Recently, we demonstrated that voxel activity patterns in visual cortex are recoded into a line-like spatial coding scheme when subjects are asked to remember and estimate the orientation of a grating or direction of moving dots^43^. As a summary statistic of each stimulus type, a line is both efficient and useful when behaviorally estimating memorized angles. These findings together with our current findings indicate that WM representations are surprisingly flexible. Within early visual cortex, sensory representations appear to transform into representations that depend on the type of memory-guided behavior. By implementing different task demands under the same sensory inputs, future studies can investigate how the dynamics of WM neural code are affected by the level of abstraction or reformation of the perceptual inputs required in memory-guided behavior.

When quantifying the polar angle response function evoked by the WM target, the widths of the functions in early visual cortex widened over time once it reached a dip at the early time points (Fig. 7), and in contrast, the width was constant throughout the delay in higher-level cortical regions. These results suggest that WM-related activation in early visual cortex may rely on feedback signals originating from the downstream regions. During target encoding, polar angle response function in early visual cortex was narrow, owing to the small receptive field sizes in these areas. Feedback signals originating from higher-level cortical areas, with larger receptive fields, might broaden the widths of the response functions in early visual cortex during WM. A shift from bottom-up sensory signals to top-down WM-related feedback may contribute to the dynamics in early visual cortex. These results are consistent with laminar recording in macaque V1^62^ and fMRI measurements of human V1^35,43,63^ showing that the persistent activity during WM in V1 has top-down origins. Beyond WM, two recent studies also employed top-down feedback signals to explain the decline of the precision of spatial representations in early visual cortex during mental imagery^64^ and episodic memory, relative to stimulus encoding.

## METHODS

### Procedures

The details of the main experiment, the passive viewing experiment and the retinotopic mapping sessions have been previously reported in^41^. In the main experiment, participants performed a memory-guided saccade task in the fMRI scanner. Each trial started with the onset of the WM target, a light gray dot (0.65° diameter) with a duration of 500 ms. The target was at 12° eccentricity and the polar angle of the target was pseudo-randomly sampled from 1 of 32 locations evenly tiling a full circle. The target was followed by a 12-second delay period, during which the participants were required to maintain their gaze at the fixation point while remembering the location of the target dot. After the delay, the fixation point changed from a light gray circle into a gray filled dot, serving as the response cue. In addition, an black ring whose radius matches the eccentricity of the target was presented. Upon the onset of the response cue, participants reported the location of the target by making a saccadic eye movement onto the black ring. The reported location was first read out by the eye tracker, and a dot was presented at the reported location. Participants were allowed to further use a manual dial to adjust the reported location, and they pressed a button to finalize the memory report. Upon the button press, an arc centered at the reported location appeared on the ring. The participants were asked to use the manual dial to adjust the length of the arc in a post-estimation wager, in which they should reflect the uncertainty of their WM by the length of the arc, the longer the arc the more uncertain. Participants finalized the uncertainty report by a button press, after which a white dot was presented at the true target location as the feedback. Participants could earn points if the true target location fell within the arc (see details in Li et al., 2021). The results of uncertainty reports are detailed in an earlier study^41^.

A subset of participants (n=6) joined an additional passive viewing experiment. The timing of this experiment was similar to the main experiment. Instead of a brief and dim WM target stimulus, we presented a salient high-contrast flickering checkerboard (0.875 deg radius; 1 cycle/deg spatial frequency; 8 Hz flicker) at the same locations as the main experiment. The checkerboard was presented for 12.5 seconds (throughout the WM target period and the delay in Figure 1A), during which the fixation point, a ‘+’ symbol, changed its weight-height ratio, and the participants were asked to attend to the fixation and detect the changes by button presses.

Each participant was scanned for a 1.5-2 hour retinotopic mapping session. The procedures of the retinotopic mapping session followed those used in Mackey et al^45^. In short, during the retinotopic mapping session, participants maintained their gaze at the fixation point while a bar sweeping across the screen 12 times per run in various directions. Participants were required to attentively track the bar and perform a motion discrimination task based on the random dot kinematograms presented within the bar apertures see details in^45^. We fit a pRF model with compressive spatial summation to the BOLD time series of the retinotopic mapping session^44,65^. We projected the voxels’ preferred phase angle and eccentricity on the cortical surface and defined ROIs by visual inspection (primarily based on the reversal of voxels’ preferred phase angle). We define bilateral dorsal visual ROIs V1, V2, V3, V3AB, IPS0, IPS1, IPS2, IPS3 and two frontal ROIs, iPCS and sPCS.

### Setup and eye tracking

Visual stimuli were presented by an LCD (VPixx ProPix) projector located behind the scanner bore. Participants viewed the stimuli through an angled mirror with a field of view of 52° by 31°. We presented a gray circular aperture (30° diameter) on the screen throughout the experiments. Eye position was recorded with a sampling rate at 500 Hz using an EyeLink 1000 Plus infrared video-based eye tracker (SR Research) mounted inside the scanner bore. We monitored gaze data and adjusted pupil/corneal reflection detection parameters as necessary during and/or between each run.

### Behavioral data analysis

As participants were allowed to manually adjust the reported location after the saccades, the final dot location after the manual adjustment was used as the participants’ memory report. Eye position was analyzed offline. The raw eye position data were first smoothed with a Gaussian kernel, and was converted into eye velocity using the eye positions of the five neighboring time points. Saccades were detected when the eye velocities exceeded the median velocity by 5 SDs with a minimum duration of 8 ms. Trials with ill-defined primary saccade, or with the magnitude of memory error or uncertainty report (arc length) larger than mean 3 standard deviations were excluded from analyses.

### Temporal generalization

We decoded the location (polar angle) of the target from the BOLD response. We focused on the BOLD response measured from 0 to 13.5 second from the delay onset (18 TRs in total). Here, decoding is a regression problem where we aimed to predict the target location *y* from single-trial BOLD response *X*(an *n_trial_* × *n_voxel_* matrix) by estimating weights*w*. As polar angle is a circular variable, we trained two regressions to predict *y_sin_* = *sin*(*y*) and *y_cos_* = *cos*(*y*) and the predicted target location was computed as *ŷ* = *atan*2(*ŷ_sin_, ŷ_cos_*). We used support vector regression (with linear kernel) in the scikit-learn Python library to estimate the weights. In a 10 fold cross-validation procedure, all the trials from a subject were separated as the training set (9/10 of the trials) and the testing set (1/10 of the trials). Regression weights were estimated using the training set’s BOLD response of a particular time point. The weights were then applied to predict the target location using the testing trials’ BOLD response. Note that the testing was not only applied to the time point same as the time point of the training data, but was applied to all the time points of the data of the testing trials to investigate the generalizability of the decoders. The performance of the decoder was quantified as (the absolute value of) decoding errors averaged across all the trials for each participant. In FIgure 1C, we report the mean decoding error averaged over participants.

### Statistical tests for the stable and dynamic code

The stable code was represented by above-chance decoding even when the training and the testing data were from different time points. Thereby we defined the stable code as the elements in the cross-decoding matrix where the decoding error was smaller than 90° (the mean decoding error under the null hypothesis that the decoder generates random predictions uniformly distributed across the entire polar angle space). We first applied t-test for each element in the cross-decoding matrix, and the t-scores were subject to cluster-based permutation test for identifying clusters exhibiting above-chance decoding performance. The permutation test was done by randomly permuting the decoders’ predicted target location and computing the t-scores summed over the element in the most-significant cluster. This procedure was repeated 1000 times resulting in a null-hypothesis distribution of the summed t-scores, which was used to decide whether a cluster was statistically significant in the cross-decoding matrix see details in^66^. The dynamic code was defined as the off-diagonal elements whose decoding performance was significantly poorer than their corresponding on-diagonal elements. Note that for an off-diagonal element to be included in a cluster, its decoding performance had to be lower than both of its corresponding on-diagonal elements. The cluster-based permutation test was done by randomly permuting the locations of the on- and off-diagonal elements. Overall, the definitions and the statistical tests for the stable and dynamic code were similar to those used in a previous monkey neurophysiological study^18^.

### Stable and dynamic subspaces

We used PCA (principal component analysis) to define low-dimensional subspaces that encode target locations. We defined a data matrix X with a size of *n_stimulus_* by *n_voxel_*, which represented the voxel activity patterns averaged across all the trials from the same stimulus location. For all the PCA conducted in this study, for the purpose of higher signal-to-noise ratio, we binned four neighboring target locations together resulting in 8 stimulus locations for the data matrix (*n_stimulus_* = 8). *n_voxel_* was the number of voxels of each ROI. X had column-wise zero empirical mean, as the mean of each column was removed. When visualizing the subspaces (Figure 2, 3E and 3G), for each ROI, we concatenated the voxels across all participants, and applied the PCA on the participant-aggregated ROI. When computing the indices that quantified the stability of the subspaces—the principal angle and the ratio of variance explained—PCA was applied to individuals’ ROIs and the indices were computed for each participant.

For the stable subspaces, we disregarded the time-varying information by averaging the data across all the time points that fell within the stable cluster during the delay (pink dashed lines in Figure 1C). Overall, all the time points within the delay were included except the first 2 or 3 TRs depending on the ROI. To obtain the principal components (PCs), we applied eigendecomposition on the covariance matrix **X**^T^**X WΛW^T^**, where each column of **W** was a unit-length vector, with a size of *n_voxel_* by 1, representing the weights of each PC and Λ was a diagonal matrix containing the corresponding eigenvalues. Throughout this study, we used the first two PCs with the largest eigenvalues to define the subspaces; thereby we focused on the weight matrix **W** with a size of *n_boxel_* by 2. To visualize the dynamics of population neural responses in the stable subspace, we projected the data of each time point into the stable subspace by computing **T = XW**, where **X** was the data matrix of a single time point and **T**, with a size of 8 by 2, was the projection of the voxel activity pattern of each stimulus location in the subspace defined by the top two PCs, PC1 and PC2. **T** is often referred to as PC scores in the context of PCA.

To investigate the dynamical aspect of the neural subspaces, we binned the BOLD response during the delay into three time windows: early, middle and late time windows, each with 5 TRs (Fig. 3A and 3B). We then estimated the subspace for each time window by applying PCA to the data matrices, **X_e_, X_m_** and **X_l_**, which were the BOLD response averaged over each of the time windows.

To quantify how the early and the late subspaces oriented in the high-dimensional neural space, we computed the principal angle, which measured the alignment between two subspaces^52–54^. The principal angle was computed using the method proposed by Björck and Golub^52^: We applied singular decomposition to the inner-product matrix 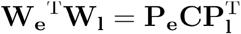, where **W_e_** and **W_l_**, both had a size of *n_voxel_* by 2, were the weighting matrices of the early and the late subspaces obtained by PCA. The matrix **C** was a diagonal matrix whose diagonal elements were the ranked (from small to large) cosines of the principal angles *θ*_1_ and *θ*_2_ **C** = diag(cos(*θ*_1_), cos(*θ*_1_). The first principal angle was reported in Figure 3B. We also computed the principal angle for two subspaces from the same time window (dashed lines in Figure 3B). This was achieved by a bootstrapping procedure with 1000 iterations. In each iteration, we resampled the trials twice, computed two subspaces using the data of the same time window and calculated the principal angle between them, resulting in a bootstrapped distribution of the principal angle within the early (or the late) time window. In addition to the principal angle, we further computed the ratio of variance explained (RVE), which quantified how much the variance explained decreased when the data of a time window was projected to the subspace of a different time window. For example, for the data of the early time window X_e_, the RVE was computed as Var(**X_e_W_l_**)/Var(**X_e_W_e_**) (Supplementary Fig. 2).

### Visualizations of WM representations

We used voxels’ pRF parameters to visualize how neural populations represent remembered locations. To compute the activation maps (Fig. 4 to Fig. 6), a grid was positioned in the visual field, centered at the fixation. The grid points evenly sampled the visual space with a step of 0.5° and the entire grid covered ±18° in both horizontal and vertical directions from the fixation point. We computed the neural activity *α_i_* for the grid point *i*, whose coordinate in the visual field was (*x_i_, y_i_*) as the weighted sum of voxel responses 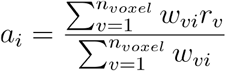. Here, *r_v_*, was the response of voxel *v*. All the voxels within an ROI were included except the voxels whose responses in the retinotopic mapping session can not be well-fitted by the pRF model (thresholding at variance explained by the pRF model = 10%). *w_vi_* was the weight of voxel *r* at grid point *i*, which was determined by the density function of a bivariate circular gaussian distribution *N*(*x_i_,y_i_*; **u**, *σ*^2^**I**), in which **u** was the voxel’s pRF center and *σ* was the voxel’s pRF size. To compute an activation map for a time window and an ROI, we did the following steps: We computed an activation map for each trial, rotated the map of each trial to align the target at 0° polar angle, averaged map over all trials for each participant, and lastly we subtracted the grand mean from the activation map.

We computed each ROI’s polar angle response function from the activation maps. The cartesian coordinates (*x_i_, y_i_*) of each grid point were first converted to phase angle *θ_i_* and eccentricity *r_i_*. We then binned the grid points based on their phase angle ranging from −180° to +180° with a step of 8°. We included the grid points with eccentricity smaller than 15°. The polar angle tuning function was computed as the activity averaged over the grid points within each bin. To quantify the dynamics of polar angle response functions, the tuning function of each time point was fitted by von Mises distributions 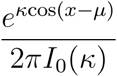, where *I*_0_(*k*) was the modified Bassel function of order 0. We reported the gain of the tuning function defined as the difference between the maximum and the minimum value of the best-fit tuning function, and the tuning width represented by the fitted *κ* value, converted to have a unit in polar angle degree (Fig. 7 and Supplementary Fig. 9).

## Acknowledgements

We thank New York University’s Center for Brain Imaging for technical support. This research was supported by the National Eye Institute (R01 EY-016407, R01 EY-027925, and R01 EY-033925 to C.E. Curtis). Hsin-Hung Li was supported by the Swartz Foundation Postdoctoral Fellowship.

---

**Supplementary Fig. 1:**
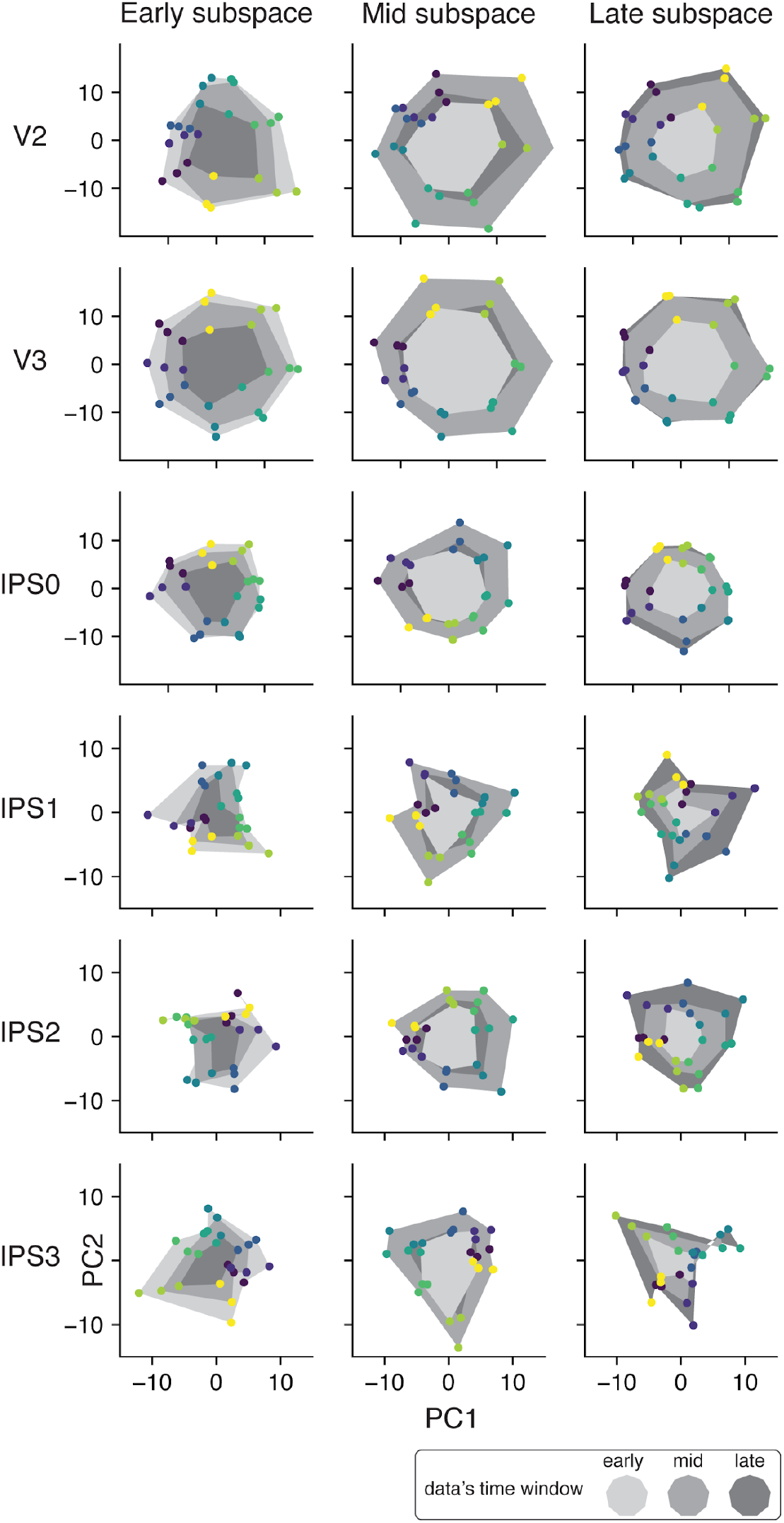
Projections of the early, mid and late (data points connected with areas with three different gray levels) voxel activity patterns of each ROI into the early, mid or late subspaces (from the late to right panel).

**Supplementary Fig. 2:**
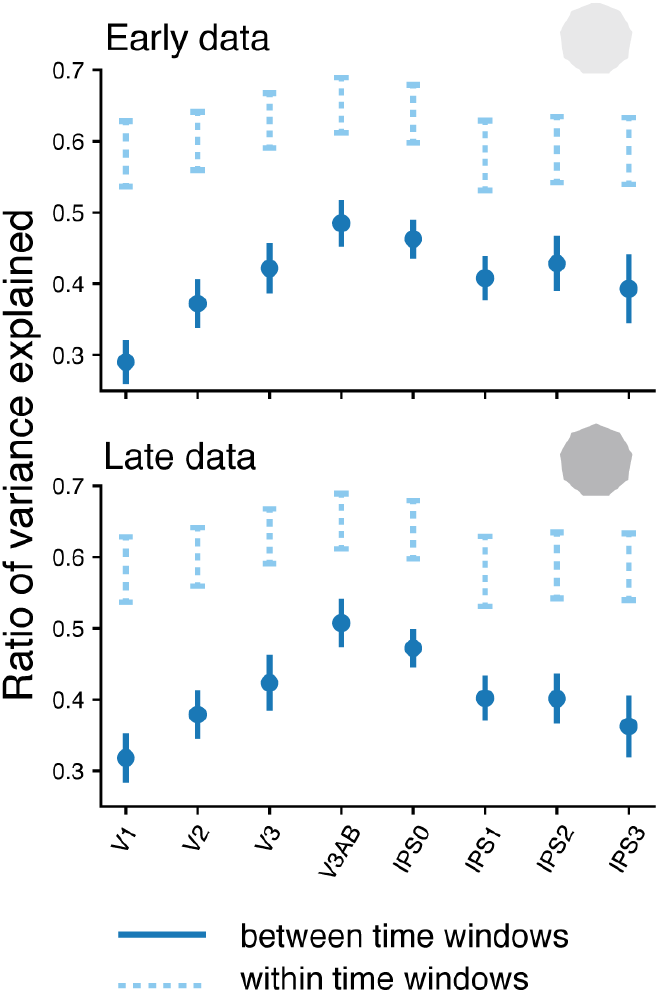
Ratio of variance explained. To quantify the reduction of the variance explained when the data of the early time window was projected into the late subspace (or vice versa), we computed the ratio between the variances explained by each of the two subspaces (RVE, dark blue data points; see Methods). We also computed the RVE within time windows by resampling the trials and computing the subspace twice using the data from the same time window (dashed light blue lines represent the 95% confidence intervals of the baseline values). The RVEs computed across time windows were smaller than the RVEs within the time windows indicating a significant reduction of variance explained when the data of one time window were projected to the subspace of the other time window. Similar to the principal angle, the (across-time) RVE varied across ROIs for both the data of the early (main effect for ROI, F(7, 91) = 2.25, p < .05) and the late time window (main effect for ROI, F(7, 91) = 2.98, p < .01).

**Supplementary Fig. 3:**
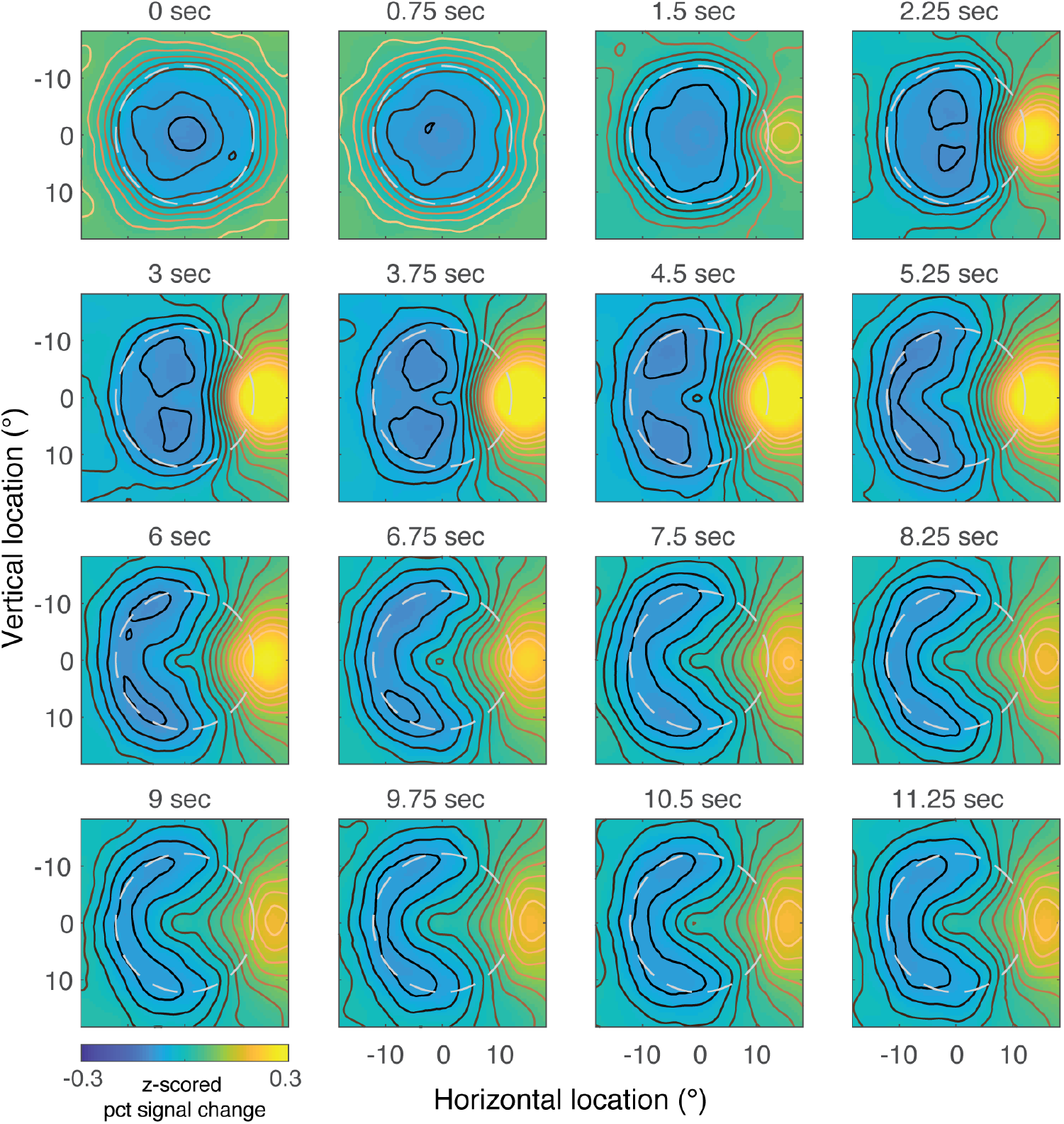
Visualization of WM dynamics in V2.

**Supplementary Fig. 4:**
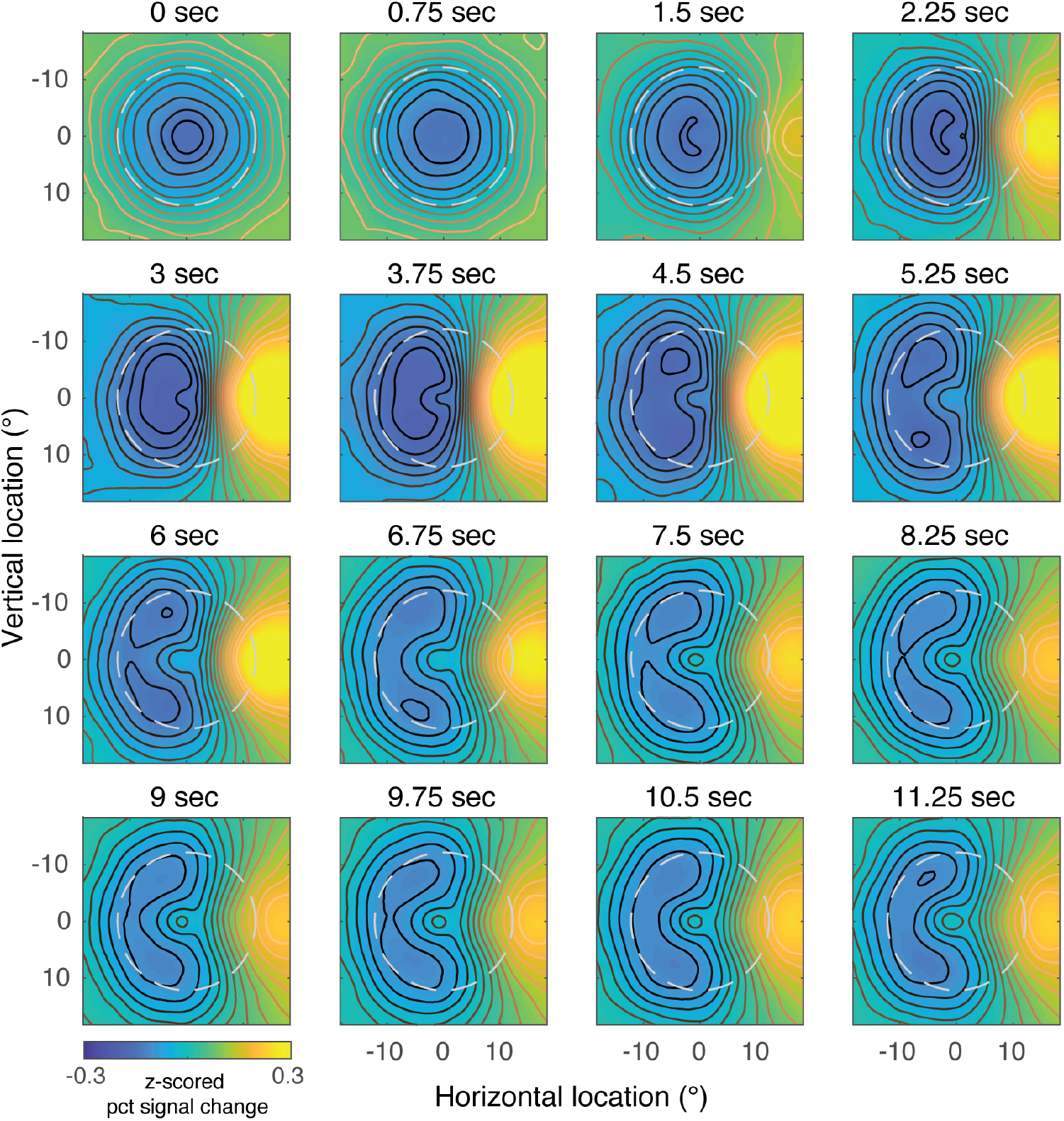
Visualization of WM dynamics in V3.

**Supplementary Fig. 5:**
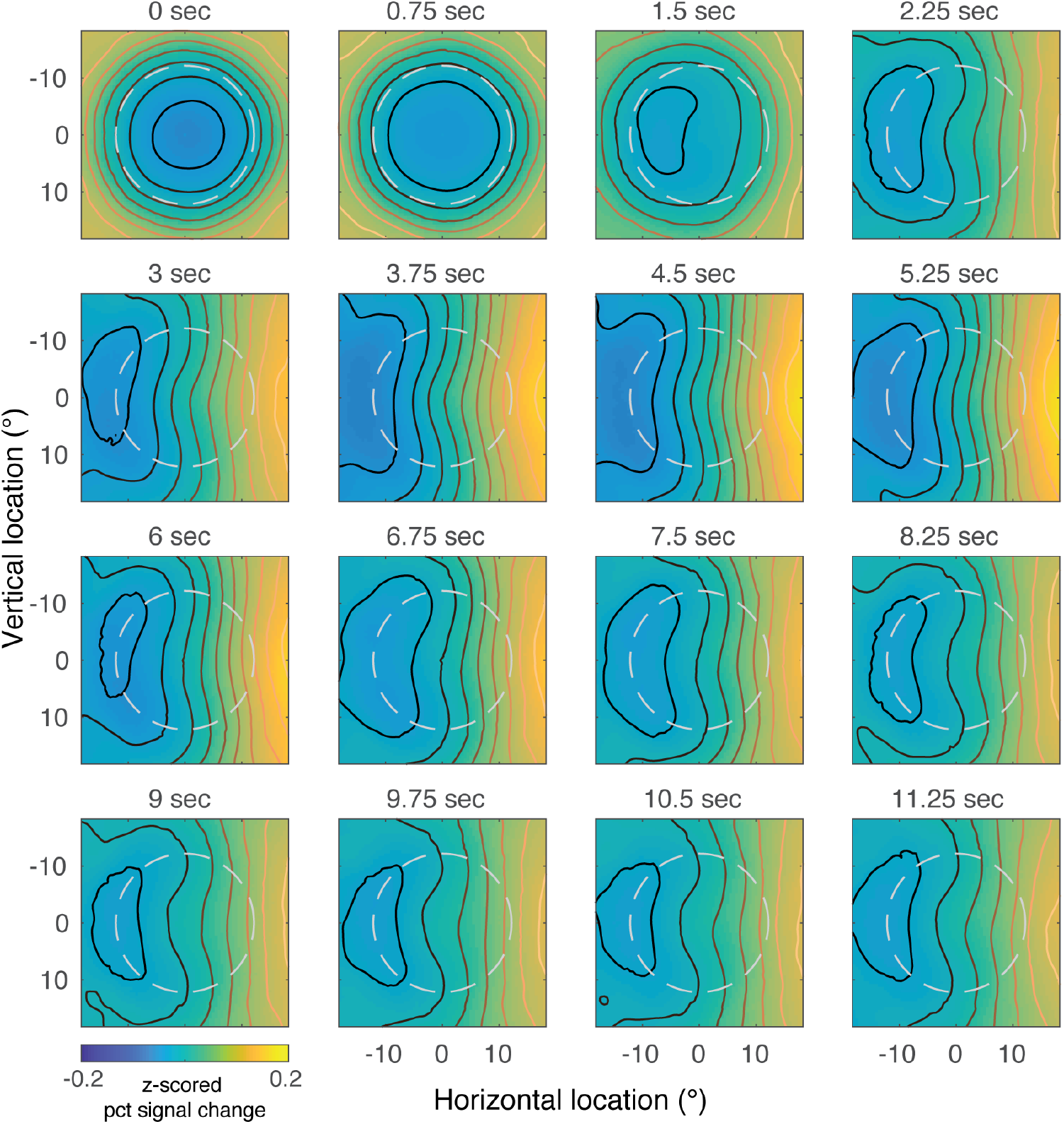
Visualization of WM dynamics in IPS0.

**Supplementary Fig. 6:**
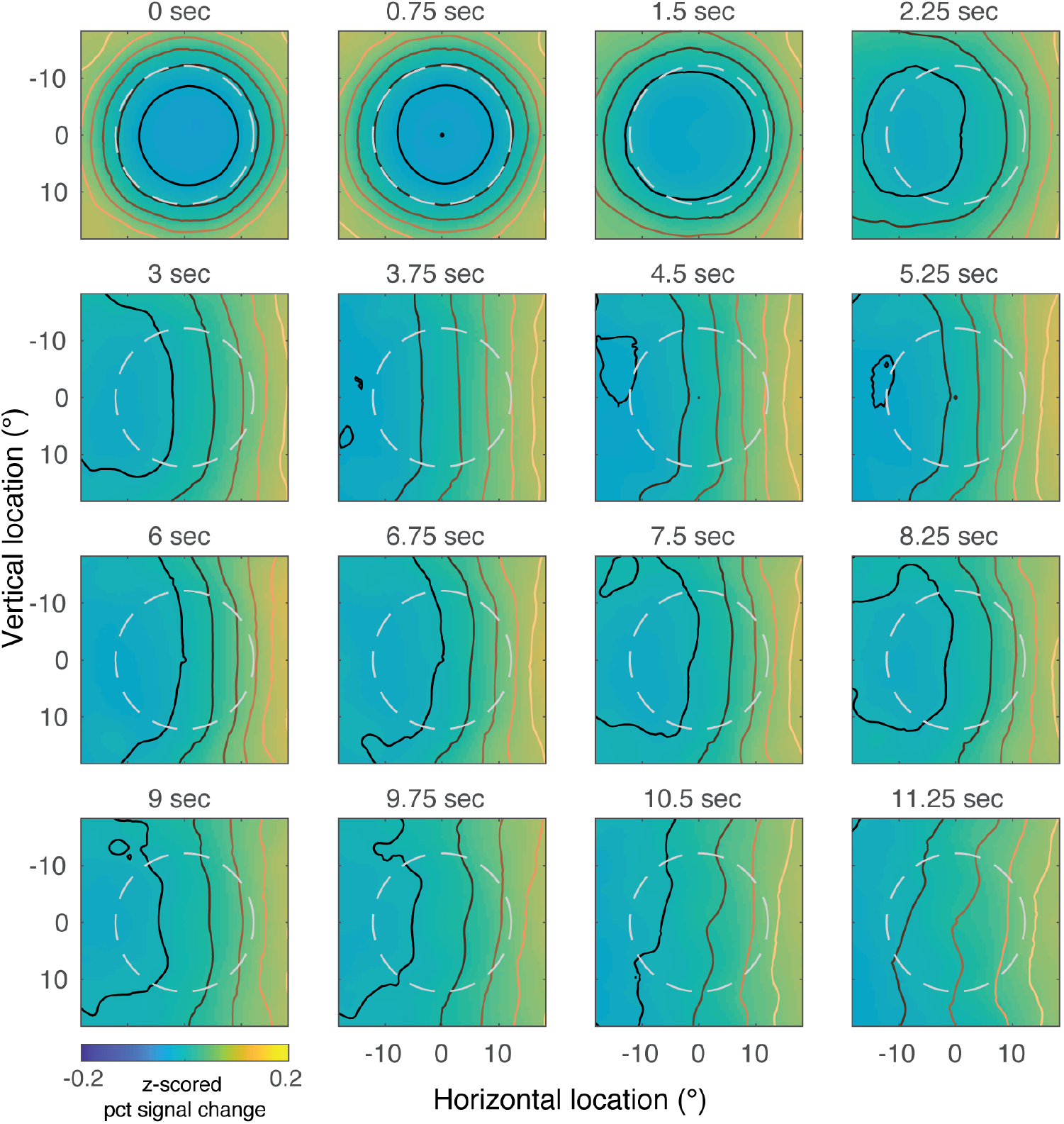
Visualization of WM dynamics in IPS1.

**Supplementary Fig. 7:**
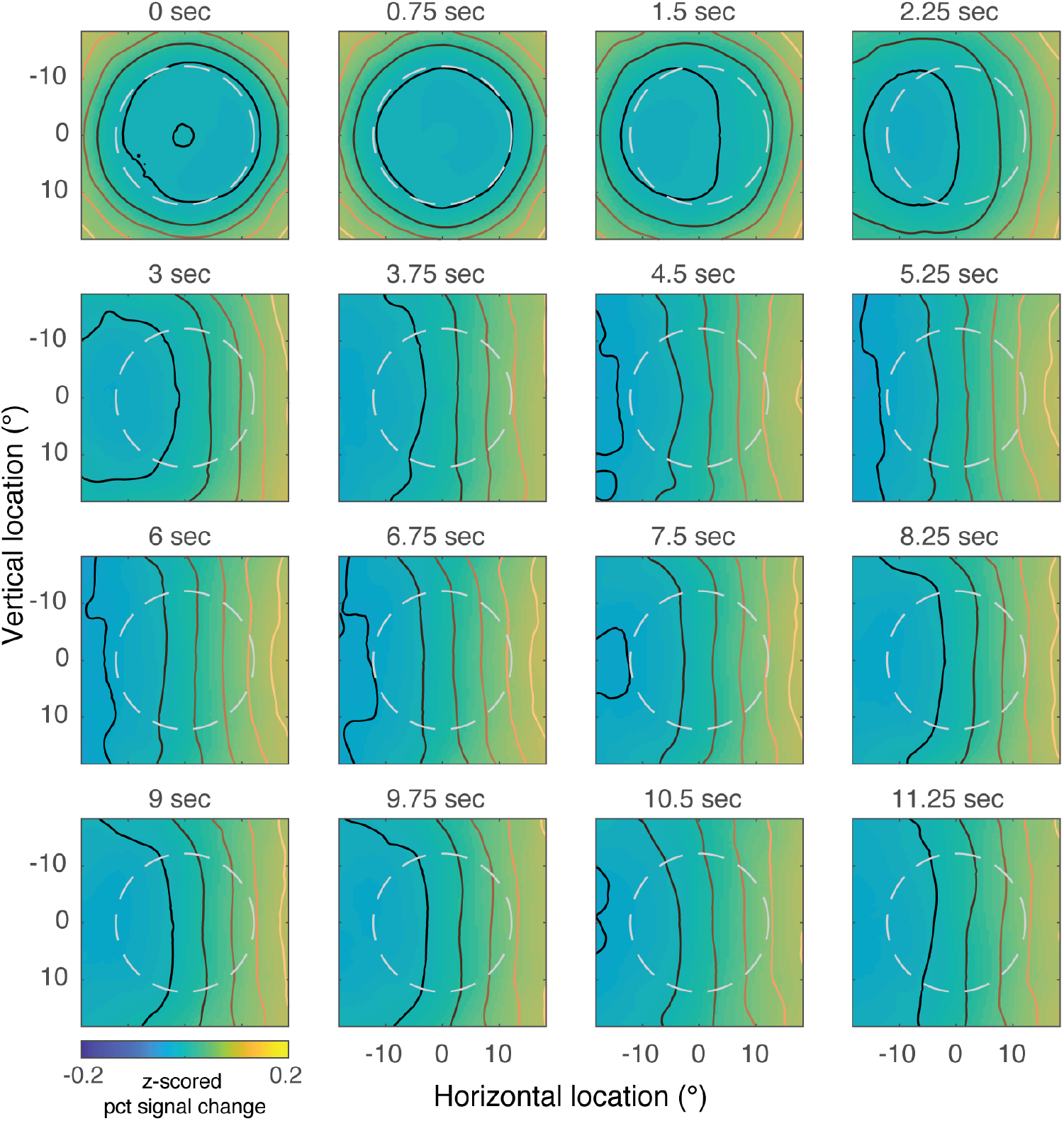
Visualization of WM dynamics in IPS2.

**Supplementary Fig. 8:**
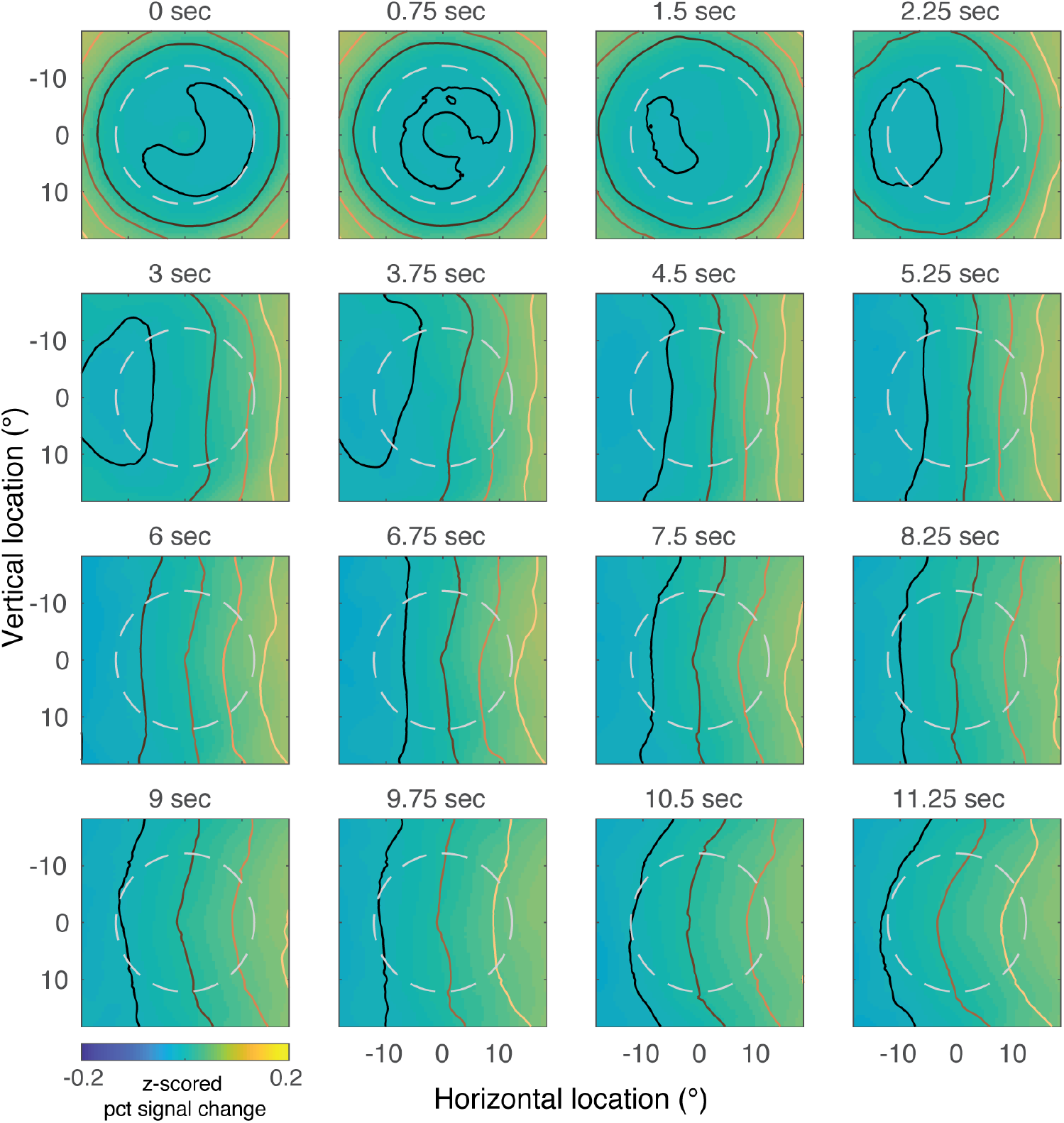
Visualization of WM dynamics in IPS3.

**Supplementary Fig. 9:**
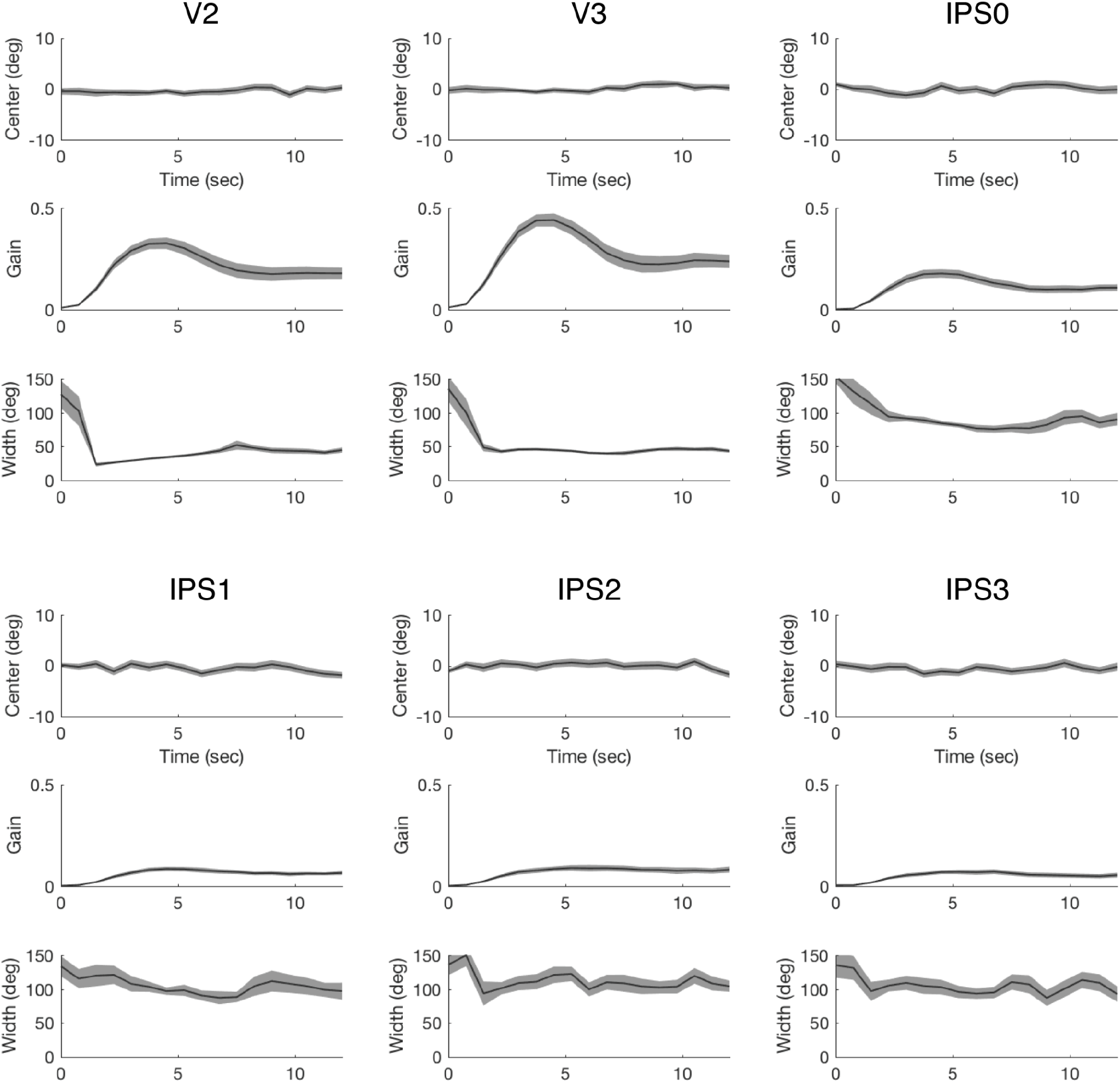
Parameter fits of the polar angle response functions. For each ROI, the fitted center, gain and width are plotted as a function of time (from delay onset). Data represents mean ±s.e.m.

## References

1. Kubota, K. & Niki, H. Prefrontal cortical unit activity and delayed alternation performance in monkeys. J. Neurophysiol. 34, 337–347 (1971).

2. Fuster, J. M. & Alexander, G. E. Neuron Activity Related to Short-Term Memory. Science vol. 173 652–654 Preprint at https://doi.org/10.1126/science.173.3997.652 (1971).

3. Curtis, C. E. & Sprague, T. C. Persistent Activity During Working Memory From Front to Back. Front. Neural Circuits 15, 696060 (2021).

4. Funahashi, S., Bruce, C. J. & Goldman-Rakic, P. S. Mnemonic coding of visual space in the monkey’s dorsolateral prefrontal cortex. J. Neurophysiol. 61, 331–349 (1989).

5. Miller, E. K., Erickson, C. A. & Desimone, R. Neural mechanisms of visual working memory in prefrontal cortex of the macaque. J. Neurosci. 16, 5154–5167 (1996).

6. Funahashi, S., Bruce, C. J. & Goldman-Rakic, P. S. Dorsolateral prefrontal lesions and *oculomotor delayed-response performance: evidence for mnemonic ‘scotomas’*. J. Neurosci. 13, 1479–1497 (1993).

7. Funahashi, S., Chafee, M. V. & Goldman-Rakic, P. S. Prefrontal neuronal activity in rhesus monkeys performing a delayed anti-saccade task. Nature 365, 753–756 (1993).

8. Romo, R., Brody, C. D., Hernández, A. & Lemus, L. Neuronal correlates of parametric working memory in the prefrontal cortex. Nature 399, 470–473 (1999).

9. Constantinidis, C., Franowicz, M. N. & Goldman-Rakic, P. S. The sensory nature of mnemonic representation in the primate prefrontal cortex. Nat. Neurosci. 4, 311–316 (2001).

10. Wasmuht, D. F., Spaak, E., Buschman, T. J., Miller, E. K. & Stokes, M. G. Intrinsic neuronal dynamics predict distinct functional roles during working memory. Nat. Commun. 9, 3499 (2018).

11. Panichello, M. F. & Buschman, T. J. Shared mechanisms underlie the control of working memory and attention. Nature 592, 601–605 (2021).

12. Brody, C. D., Hernández, A., Zainos, A. & Romo, R. Timing and neural encoding of *somatosensory parametric working memory in macaque prefrontal cortex*. Cereb. Cortex 13, 1196–1207 (2003).

13. Baeg, E. H. et al. Dynamics of population code for working memory in the prefrontal cortex. Neuron 40, 177–188 (2003).

14. Shafi, M. et al. Variability in neuronal activity in primate cortex during working memory tasks. Neuroscience 146, 1082–1108 (2007).

15. Barak, O., Tsodyks, M. & Romo, R. Neuronal population coding of parametric working memory. J. Neurosci. 30, 9424–9430 (2010).

16. Harvey, C. D., Coen, P. & Tank, D. W. Choice-specific sequences in parietal cortex during a virtual-navigation decision task. Nature 484, 62–68 (2012).

17. Murray, J. D. et al. Stable population coding for working memory coexists with heterogeneous neural dynamics in prefrontal cortex. Proc. Natl. Acad. Sci. U. S. A. 114, 394–399 (2017).

18. Spaak, E., Watanabe, K., Funahashi, S. & Stokes, M. G. Stable and Dynamic Coding for Working Memory in Primate Prefrontal Cortex. The Journal of Neuroscience vol. 37 6503–6516 Preprint at https://doi.org/10.1523/jneurosci.3364-16.2017 (2017).

19. Wolff, M. J., Ding, J., Myers, N. E. & Stokes, M. G. Revealing hidden states in visual working memory using electroencephalography. Front. Syst. Neurosci. 9, 123 (2015).

20. Wolff, M. J., Jochim, J., Akyürek, E. G., Buschman, T. J. & Stokes, M. G. Drifting codes within a stable coding scheme for working memory. PLoS Biol. 18, e3000625 (2020).

21. Serences, J. T., Ester, E. F., Vogel, E. K. & Awh, E. Stimulus-specific delay activity in human primary visual cortex. Psychol. Sci. 20, 207–214 (2009).

22. Harrison, S. A. & Tong, F. Decoding reveals the contents of visual working memory in early visual areas. Nature 458, 632–635 (2009).

23. Jerde, T. A., Merriam, E. P., Riggall, A. C., Hedges, J. H. & Curtis, C. E. Prioritized maps of space in human frontoparietal cortex. J. Neurosci. 32, 17382–17390 (2012).

24. Riggall, A. C. & Postle, B. R. The relationship between working memory storage and elevated activity as measured with functional magnetic resonance imaging. J. Neurosci. 32, 12990–12998 (2012).

25. Emrich, S. M., Riggall, A. C., Larocque, J. J. & Postle, B. R. Distributed patterns of activity in sensory cortex reflect the precision of multiple items maintained in visual short-term memory. J. Neurosci. 33, 6516–6523 (2013).

26. Ester, E. F., Anderson, D. E., Serences, J. T. & Awh, E. A neural measure of precision in visual working memory. J. Cogn. Neurosci. 25, 754–761 (2013).

27. Lee, S.-H., Kravitz, D. J. & Baker, C. I. Goal-dependent dissociation of visual and prefrontal cortices during working memory. Nat. Neurosci. 16, 997–999 (2013).

28. Xing, Y., Ledgeway, T., McGraw, P. V. & Schluppeck, D. Decoding working memory of stimulus contrast in early visual cortex. J. Neurosci. 33, 10301–10311 (2013).

29. Sprague, T. C., Ester, E. F. & Serences, J. T. Reconstructions of information in visual spatial working memory degrade with memory load. Curr. Biol. 24, 2174–2180 (2014).

30. Sreenivasan, K. K., Vytlacil, J. & D’Esposito, M. Distributed and dynamic storage of working memory stimulus information in extrastriate cortex. J. Cogn. Neurosci. 26, 1141–1153 (2014).

31. Ester, E. F., Sprague, T. C. & Serences, J. T. Parietal and Frontal Cortex Encode Stimulus-Specific Mnemonic Representations during Visual Working Memory. Neuron 87, 893–905 (2015).

32. Sprague, T. C., Ester, E. F. & Serences, J. T. Restoring Latent Visual Working Memory Representations in Human Cortex. Neuron 91, 694–707 (2016).

33. Bettencourt, K. C. & Xu, Y. Decoding the content of visual short-term memory under distraction in occipital and parietal areas. Nat. Neurosci. 19, 150–157 (2016).

34. Christophel, T. B., Klink, P. C., Spitzer, B., Roelfsema, P. R. & Haynes, J.-D. The Distributed Nature of Working Memory. Trends Cogn. Sci. 21, 111–124 (2017).

35. Rahmati, M., Saber, G. T. & Curtis, C. E. Population Dynamics of Early Visual Cortex during Working Memory. J. Cogn. Neurosci. 30, 219–233 (2018).

36. Christophel, T. B., Iamshchinina, P., Yan, C., Allefeld, C. & Haynes, J.-D. Cortical specialization for attended versus unattended working memory. Nat. Neurosci. 21, 494–496 (2018).

37. Lorenc, E. S., Sreenivasan, K. K., Nee, D. E., Vandenbroucke, A. R. E. & D’Esposito, M. Flexible Coding of Visual Working Memory Representations during Distraction. J. Neurosci. 38, 5267–5276 (2018).

38. Rahmati, M., DeSimone, K., Curtis, C. E. & Sreenivasan, K. K. Spatially Specific Working Memory Activity in the Human Superior Colliculus. J. Neurosci. 40, 9487–9495 (2020).

39. Brissenden, J. A., Tobyne, S. M., Halko, M. A. & Somers, D. C. Stimulus-Specific Visual Working Memory Representations in Human Cerebellar Lobule VIIb/VIIIa. J. Neurosci. 41, 1033–1045 (2021).

40. Hallenbeck, G. E., Sprague, T. C., Rahmati, M., Sreenivasan, K. K. & Curtis, C. E. Working memory representations in visual cortex mediate distraction effects. Nat. Commun. 12, 4717 (2021).

41. Li, H.-H., Sprague, T. C., Yoo, A. H., Ma, W. J. & Curtis, C. E. Joint representation of working memory and uncertainty in human cortex. Neuron 109, 3699–3712.e6 (2021).

42. Rademaker, R. L., Chunharas, C. & Serences, J. T. Coexisting representations of sensory and mnemonic information in human visual cortex. Nat. Neurosci. 22, 1336–1344 (2019).

43. Kwak, Y. & Curtis, C. E. Unveiling the abstract format of mnemonic representations. Neuron (2022) doi:10.1016/j.neuron.2022.03.016.

44. Dumoulin, S. O. & Wandell, B. A. Population receptive field estimates in human visual cortex. Neuroimage 39, 647–660 (2008).

45. Mackey, W. E., Winawer, J. & Curtis, C. E. Visual field map clusters in human frontoparietal cortex. Elife 6, (2017).

46. Wimmer, K., Nykamp, D. Q., Constantinidis, C. & Compte, A. Bump attractor dynamics in *prefrontal cortex explains behavioral precision in spatial working memory*. Nat. Neurosci. 17, 431–439 (2014).

47. Stokes, M. G. et al. Dynamic coding for cognitive control in prefrontal cortex. Neuron 78, 364–375 (2013).

48. King, J.-R., Pescetelli, N. & Dehaene, S. Brain Mechanisms Underlying the Brief Maintenance of Seen and Unseen Sensory Information. Neuron 92, 1122–1134 (2016).

49. Libby, A. & Buschman, T. J. Rotational dynamics reduce interference between sensory and memory representations. Nat. Neurosci. 24, 715–726 (2021).

50. Wan, Q., Menendez, J. A. & Postle, B. R. Priority-based transformations of stimulus representation in visual working memory. PLoS Comput. Biol. 18, e1009062 (2022).

51. Churchland, M. M. et al. Neural population dynamics during reaching. Nature 487, 51–56 (2012).

52. Björck, Áke & Golub, G. H. Numerical methods for computing angles between linear subspaces. Math. Comput. 27, 579–594 (1973).

53. Gallego, J. A. et al. Cortical population activity within a preserved neural manifold underlies multiple motor behaviors. Nat. Commun. 9, 4233 (2018).

54. Xie, Y. et al. Geometry of sequence working memory in macaque prefrontal cortex. Science 375, 632–639 (2022).

55. Zhou, Y., Curtis, C. E., Sreenivasan, K. K. & Fougnie, D. Common Neural Mechanisms Control Attention and Working Memory. J. Neurosci. 42, 7110–7120 (2022).

56. Kok, P. & de Lange, F. P. Shape perception simultaneously up-and downregulates neural activity in the primary visual cortex. Curr. Biol. 24, 1531–1535 (2014).

57. Favila, S. E., Kuhl, B. A. & Winawer, J. Perception and memory have distinct spatial tuning properties in human visual cortex. Nat. Commun. 13, 5864 (2022).

58. Wandell, B. A. & Winawer, J. Computational neuroimaging and population receptive fields. Trends Cogn. Sci. 19, 349–357 (2015).

59. Watanabe, K. & Funahashi, S. Prefrontal delay-period activity reflects the decision process of a saccade direction during a free-choice ODR task. Cereb. Cortex 17 Suppl 1, i88–100 (2007).

60. Constantinidis, C., Franowicz, M. N. & Goldman-Rakic, P. S. Coding specificity in cortical microcircuits: a multiple-electrode analysis of primate prefrontal cortex. J. Neurosci. 21, 3646–3655 (2001).

61. Libby, A. & Buschman, T. J. Rotational dynamics reduce interference between sensory and memory representations. Nat. Neurosci. 24, 715–726 (2021).

62. van Kerkoerle, T., Self, M. W. & Roelfsema, P. R. Layer-specificity in the effects of attention and working memory on activity in primary visual cortex. Nat. Commun. 8, 13804 (2017).

63. Saber, G. T., Pestilli, F. & Curtis, C. E. Saccade planning evokes topographically specific activity in the dorsal and ventral streams. J. Neurosci. 35, 245–252 (2015).

64. Breedlove, J. L., St-Yves, G., Olman, C. A. & Naselaris, T. Generative Feedback Explains Distinct Brain Activity Codes for Seen and Mental Images. Curr. Biol. 30, 2211–2224.e6 (2020).

65. Kay, K. N., Winawer, J., Mezer, A. & Wandell, B. A. Compressive spatial summation in human visual cortex. J. Neurophysiol. 110, 481–494 (2013).

66. Maris, E. & Oostenveld, R. Nonparametric statistical testing of EEG-and MEG-data. J. Neurosci. Methods 164, 177–190 (2007).

